# Use of a corneal impression membrane for the detection of Herpes Simplex Virus type-1

**DOI:** 10.1101/455550

**Authors:** Matthias Brunner, Tobi Somerville, Caroline E Corless, Jayavani Myneni, Tara Rajhbeharrysingh, Stephenie Tiew, Timothy Neal, Stephen B. Kaye

## Abstract

**Purpose:** To investigate the use of a corneal impression membrane (CIM) for the detection of Herpes Simplex Virus type 1 (HSV-1) in suspected Herpes Simplex Keratitis (HSK).

**Materials and Methods:** In the laboratory study, swabs and CIMs made from polytetrafluoroethylene were spiked with different concentrations of HSV-1. DNA was extracted and real time PCR undertaken using 2 sets of primers. In the clinical study consecutive patients presenting with suspected HSK were included. For each patient, samples were collected from corneal lesions with a swab and a CIM in random order. Clinical details were collected using a standardised clinical form and patients were categorized into probable, presumed and possible HSK.

**Results:** There was no difference in the performance of both primer sets for all HSV-1 dilutions (p=0.83) or between a CIM and a swab (p=0.18). 110 patients were included. Seventy-three patients (66.4%) had probable, 20 patients (18.2%) presumed, and 17 patients (15.5%) possible HSV-1 keratitis. The HSV-1 detection rate was significantly higher using a CIM (40/110, 36.4%) than a swab (28/110, 25.5%) (p=0.004). In the probable HSV keratitis group, the detection rate using a CIM was 43.8% compared to 27.4% for a swab (p=0.004). The Cp values obtained for the conjunctival swabs were higher than those obtained for the CIMs (p<0.001).

**Conclusions:** In suspected HSK, a CIM is a useful alternative to a swab and more likely to detect the presence of HSV-1.

## Introduction

Herpes Simplex Virus Type 1 (HSV-1) keratitis is a leading cause of visual impairment.(1) The annual incidence of HSV-1 keratitis (HSK) in the United States and France has been estimated at 8.4 and 31.5 per 100,000.(2) HSK most commonly presents as an epithelial keratitis with virus replicating in, and destroying, epithelial cells.(3) The lesions start as punctuate vesicular eruptions in the corneal epithelium which coalesce into dendritiform lesions and occasionally into larger non-linear geographic lesions.(4) HSK is prone to recurrence, usually manifesting as a dendritiform keratitis and or an interstitial stromal keratititis.

Although some of the clinical features of HSK keratitis are characteristic, there are other diseases and infections with similar features such as acanthamoeba keratitis. There has also been an increase in HSV-1 resistance to topical and systemic antiviral agents.(5-7) It is important, therefore, to identify and if possible, isolate HSV-1 for clinical management. Laboratory tests are aimed at cell cytology, viral antigen detection (immunoassays), viral DNA detection (polymerase chain reaction) and virus isolation (tissue culture).(8) Cytology depends on detecting the presence of intranuclear inclusions and multinucleated giant cells. It is seldom used as it has a low specificity and sensitivity (57%).(9) Isolation of HSV-1 by culture has a low sensitivity but is the standard for diagnostic specificity, potential strain identification and epidemiological tracing.(9) Enzyme or fluorescence based immunohistochemical (IHC) techniques have good sensitivity but can be difficult to interpret. The polymerase chain reaction (PCR) carries a high sensitivity but is prone to contamination and false positives. This has been addressed in part by quantification of the amount of viral DNA present using real-time PCR, which is also useful for evaluating the efficacy of antiviral medications.(10) Real-time PCRs have been developed for the detection of HSV-1 and 2 from clinical samples including genital swabs and CSF.(11-14)

Collection of samples from corneal lesions in HSK is conventionally undertaken using a swab (cotton tipped) or less commonly a blade or a needle to scrape the edges of the ulcer. Swabs, however, are cumbersome, may be difficult to localise to the ulcer using slit lamp biomicroscopy and may come into contact with the conjunctiva and or the eyelids. It is unclear whether in clinical practice the majority of specimens are collected from the conjunctiva and tear film rather than from the ulcer itself. Sharp instruments, such as a blade or needle, more commonly used in suspected bacterial or fungal ulcers, are seldom used for the detection of HSV-1, particularly because they require expertise and may lead to further corneal injury.

Corneal impression membranes (CIM) made for example, from cellulose acetate or polytetrafluoroethylene (PTFE) have been used to collect samples from the cornea and or the conjunctiva. This method, called impression cytology (IC), has been shown to reliably remove epithelial surface cells from the ocular surface for diagnostic purposes in a variety of ocular surface disease.(15-18) As has been shown in cases of suspected bacterial, acanthamoeba and fungal keratitis, use of a CIM has several practical advantages over conventional methods using swabs or sharp instruments with good isolation rates.(16) This technique is easy to perform, less traumatic and invasive for the patient and if needed, can be sized to cover the entire ulcer.(16)

To date, there are no clinical data available on the comparison of detection rates for HSV-1 using the above mentioned collection techniques. The aim of this study was to investigate and compare *in vitro* and *in vivo*, the ability of a CIM and a swab to detect the presence of HSV-1.

## Materials and Methods

### HSV-1 PCR

The performance of two HSV-1 primers in PCRs were evaluated, Bennett et al^13^ and Dupuis et al(13, 14). Master mixes for both primers comprised Roche Lightcycler 480 Probes Master Mix (Roche, Risch-Rotkreuz, Switzerland) and oligonucleotides (Eurogentec) with amplicons detected using an FAM labelled fluorescent probe (Eurogentec). Human RNaseP gene and GAPDH oligonucleotides (Eurogentec, Liège, Belgium) were used as internal amplification controls. (19, 20) Ten microlitre (µL) aliquot of eluted nucleic acid was added to 15µL master mix in a 96 well reaction plate. The parameters using a real-time PCR LC480 analyser (Roche) were 95°C for 5 mins, 45 cycles of 95 C for 10 secs, 60 C for 45 secs and 72 C for 1 sec and a final cooling step of 40 C for 30 secs. A crossing point (Cp) value for the maximum number of cycles required for HSV-1 DNA amplification was set at less than or equal to 38.7 based on the work of Bennett et al (2013).(14)

## Laboratory Study

### Recovery of HSV-1 DNA from CIM and a swab

Sterile CIM (Biopore filter paper, diameter 4mm, pore size 0.4 µm; Millicell-CM 0.4 µm PICM 01250, Millipore Corp, Bedford, MA, USA) and cotton tipped swabs (Sigma) were used. HSV-1 virus stocks of 10^4^, 10^3^, 10^2^ and 10 genome copies/mL were made by diluting cultured virus from a clinical isolate in buffer containing detergent (Hologic Apitima, Hologic, Massachusetts, USA). The number of virus genomes (copies/mL) was determined using a commercial quantitative HSV-1/2 PCR kit (QIAGEN, Hilden, Germany). To mimic clinical samples, human genomic DNA (Roche Diagnostics, Burgess Hill, UK) was diluted 10,000-fold and 1µL added to each CIM and swab. This dilution resulted in a Cp value of 29 to 31 which was comparable to that obtained from a clinical sample.

Five µL of titrated HSV-1 cultured virus stock was applied to CIMs and allowed to soak into the material. 400µL buffer containing detergent (Hologic) was added to one set of each duplicated sample and vortexed for 5 seconds before transfer of the liquid into a secondary tube for automated DNA extraction using the Roche MagNA Pure Compact and the Nucleic Acid Isolation Kit I DNA (Roche) with an elution volume of 50µL. The CIM was left in the primary tube as it would have blocked the pipette tip on the extraction instrument if transferred. The second set of each duplicant was stored at ambient temperature for 24 hours before the addition of 400µL buffer containing detergent and DNA extraction. For comparison, simulated corneal swabs were similarly inoculated. Five µL HSV-1 at each dilution was applied to a cotton swab which was then added to a tube of 3mL Sigma Virocult viral transport medium (MWE, Wiltshire, UK) and vortexed for five seconds. A 400µL aliquot of viral transport medium was transferred to a secondary tube before nucleic acid extraction as before.

Topical anaesthetic is usually applied to the eye prior to collection of samples from the cornea. To investigate possible inhibitory effects on the PCR and HSV-1 recovery(21), after adding 5µL HSV-1 at 10^2^ or 10 virus copies/mL to each CIM, 1µL of undiluted and diluted (1 in 10^2^ and 10^3^) proxymethocaine (Bausch & Lomb UK Limited, Kingston-upon-Thamas, Surrey, UK) was added to each CIM before the addition of buffer containing detergent (Hologic) and DNA extraction.

## Clinical Study

### Patient selection

Consecutive patients presenting to The Royal Liverpool University Hospital with suspected epithelial HSK were prospectively recruited between June 2016 and December 2017. Patient demographics and clinical details including previous ophthalmic history, best-corrected visual acuity (BCVA), characteristics of lesions, extraocular manifestations and treatment, were collected using a standardised clinical form. Patients were categorised into probable, presumed and possible HSK by two independent observers. Probable HSK was defined as the presence of a dendritic or geographic ulcer with or without an associated corneal stromal keratitis. Presumed HSK was defined as an atypical keratitis (non-dendritiform or non-geographic ulcers) with or without stromal lesions in a patient with a history of a previous and or recurrent HSK. Possible HSK was defined as clinical microbial keratitis in which HSV-1 was a consideration, but for which there were no typical HSK features and no history of HSK. Patients below age 18 years, with incomplete data or no matching were excluded. All included patients provided informed consent. The study received Institutional Review Board approval from the ethical committee of The Royal Liverpool and Broadgreen University Hospital and was conducted according to the ethical standards set out in the 1964 Declaration of Helsinki, as revised in 2000.

### Sample Collection

Two samples (corneal swab and CIM) were collected from the corneal lesion at presentation. The order of collection was randomised. Following instillation of a topical anaesthetic (one drop of 0.5% proxymethacaine) to the lower conjunctival fornix, a sample was collected. The swab was rolled across the corneal lesion and placed in 3 mL Sigma Virocult viral transport medium. This was followed or preceded by application of a CIM (4mm diameter millipore filter paper, pore size 0.4 µm), to the surface of the lesion for 5 seconds using a sterilised forceps. The filter paper was then transferred to a sterile tube and transported to the laboratory for DNA extraction and PCR.

### Statistical Methods

A sample size of 100 patients was based on alpha of 0.05, sensitivity 0.85, specificity 0.90, precision 0.1 and an assumed viral detection rate of 30%-35% with corneal swabs.(22) Statistical analysis was performed using SPSS (version 22). Independent t-tests were used to compare recovery of HSV-1 DNA between CIMs extracted at 0h and 24h and between the CIMs and swabs. Chi-square tests were used to compare the differences in HSV-1 detection rate between the CIM and conjunctival swab. One-way ANOVA was used to test for differences between the Dupuis and Bennett primer Cp values for the conjunctival swabs and CIMs. Post hoc analysis was carried out using Bonferroni post hoc test.

## Results

There was no evidence of inhibition of the HSV-1 PCR using CIM inoculated with 10 and 100 HSV-1 copies/mL in the presence or absence of different concentrations of eye drops (p=0.91, Table 1).

**Table 1.**
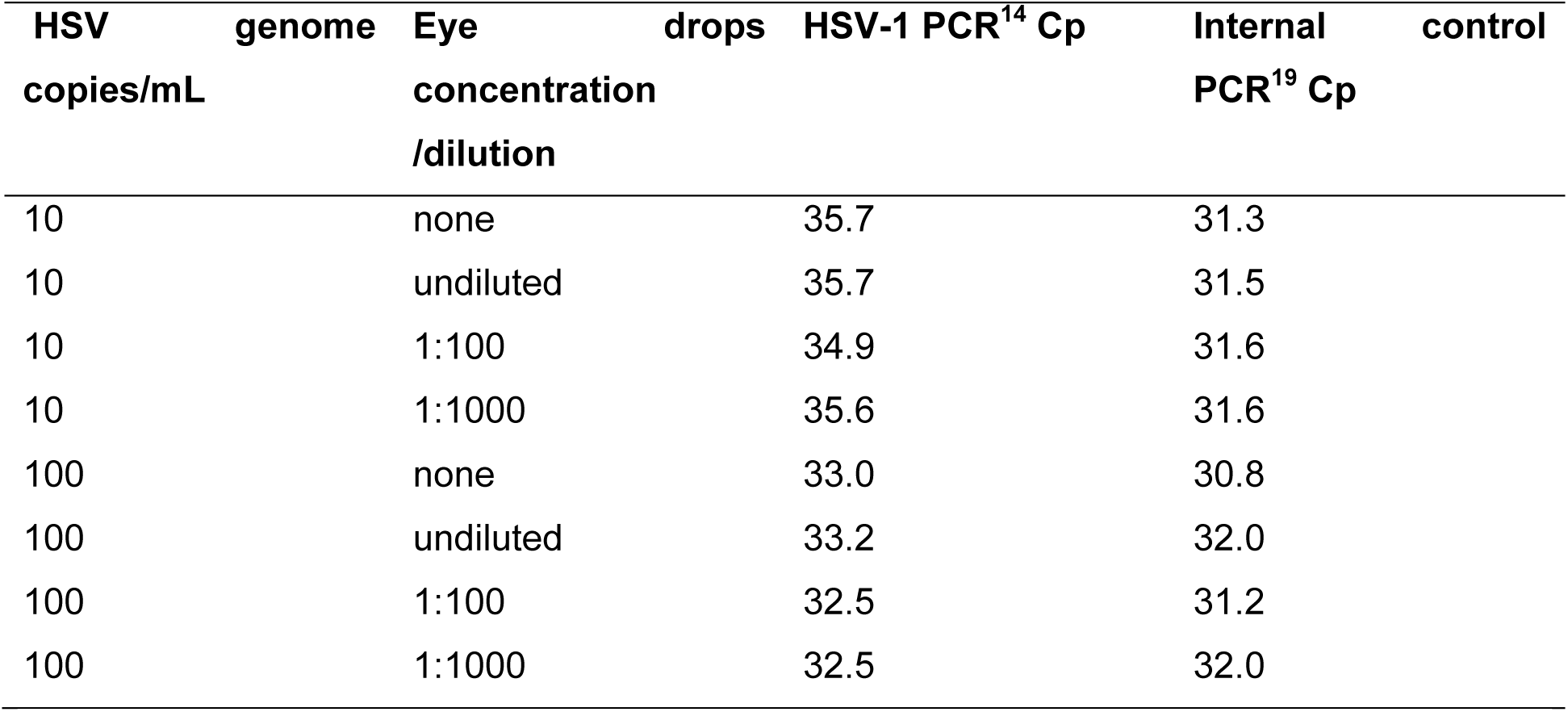
Effect of eye drop concentration on HSV-1 PCR detection

DNA extracted immediately after HSV-1 inoculation (wet) or 24 hours after HSV-1 inoculation with dry storage yielded similar Cp values for both PCRs for all HSV-1 dilutions (Figure 1). (p=0.83). The Cp PCR values obtained following inoculation with a CIM were approximately 3-PCR cycles lower than the corresponding Cp values from a swab but this was not significant (p=0.18).

**Figure 1.**
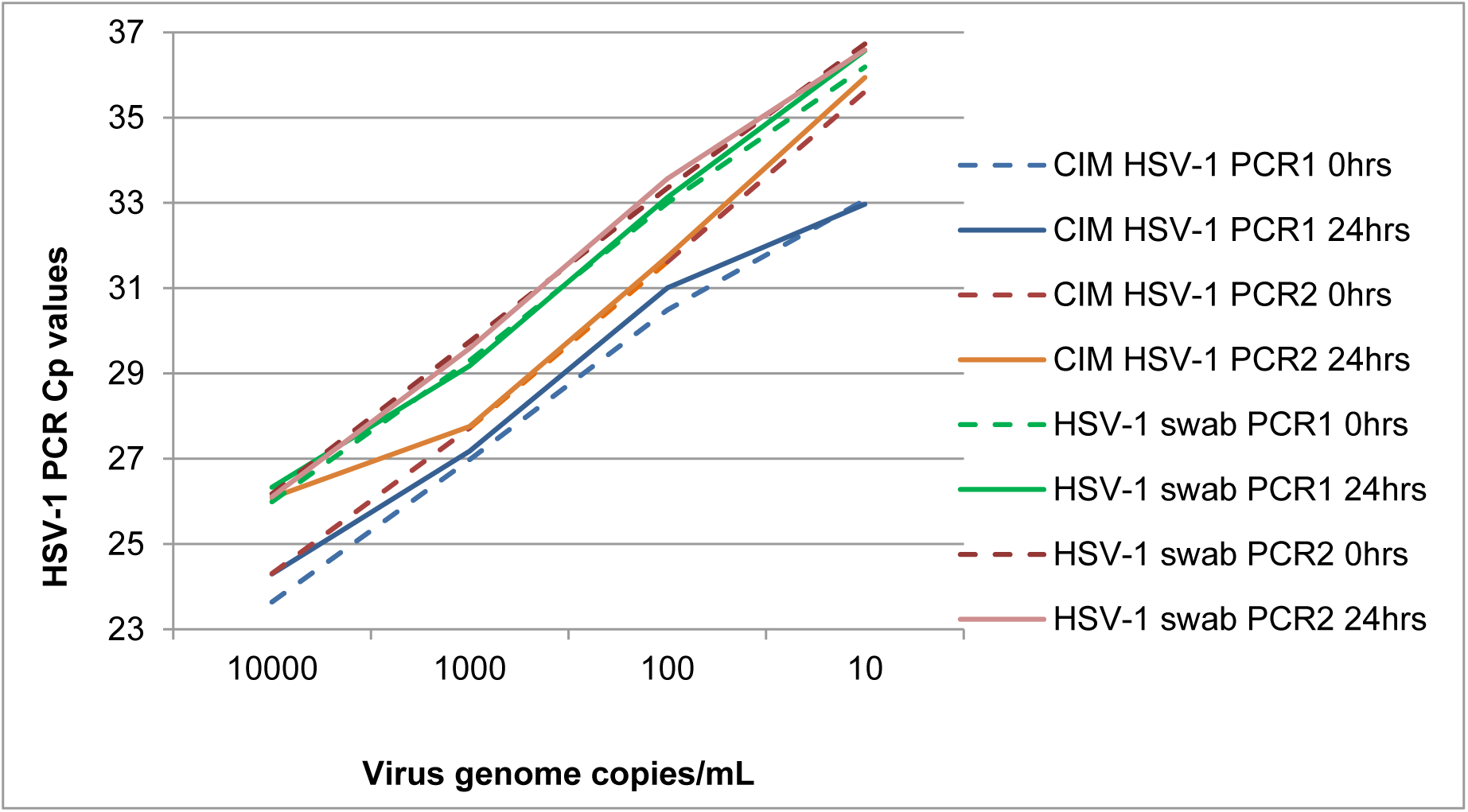
Comparative sensitivities of HSV-1 PCRs with CIM and eye swabs. Amplification of HSV-1 DNA from swabs and corneal impression membranes using Dupois and Bennet primers.(13, 14) HSV-1 DNA was extracted immediately (0hrs) and 24hrs after inoculation. Cp values plotted for serial 10-fold dilutions of virus genome copies/mL.

### Clinical study

In total, 110 consecutive patients (56 males and 54 females) were included (mean age 55.4 years, SD 17.2). As determined by two independent observers; 73 patients had probable, 20 patients presumed and 17 patients possible HSV-1 keratitis. Forty-five patients (40.9%) had a history of recurrent disease with previous episodes of HSK of whom 31 patients had pre-existing corneal scarring and 27 of these had associated corneal neovascularisation. Fifty-eight patients (52.7%) had best corrected visual acuity of worse than 6/12 at presentation.

The HSV-1 detection rate was significantly higher using a CIM (40/110, 36.4%) than a swab (28/110, 25.5%) (p=0.004). In the probable HSV keratitis group, the detection rate using a CIM was 43.8% compared to 27.4% for a swab (p=0.004). No significant difference was found between the HSV-1 detection rates between the CIM and the conjunctival swab in the presumed and possible HSK groups (Tables 2 and 3). The Cp values obtained for the conjunctival swabs were higher than those obtained for the CIMs (p<0.001, table 3). Comparing both sets of HSV-1 PCR primers, there was one sample out of 110 where the Dupuis PCR was borderline positive (37.02) and the Bennett negative. There was one sample which was inhibitory with the Dupuis PCR but was positive using the Bennett primers for both the CIM and swab. Post hoc analysis demonstrated no difference between the two sets of HSV-1 primers Cp values (table 3).

**Table 2.** HSV-1 detection rates using Dupuis real-time HSV-1 PCR

| HSK | n | Swab positive (%) | CIM positive (%) | P value |
| --- | --- | --- | --- | --- |
| Overall | 110 | 28 (25.5) | 40 (36.4) | 0.004 |
| Probable* | 73 | 20 (27.4) | 32 (43.8) | 0.004 |
| Presumed* | 20 | 6 (30.0) | 6 (30.0) | 1 |
| Possible* | 17 | 2 (11.8) | 2 (11.8) | 1 |

\*As determined by two independent observers.

**Table 3.** HSV-1 PCR Cp values using Dupuis and Bennett primers

| HSK | n | Swab Dupuis<br>Primers<br>Cp Mean (SD) | CIM Dupuis<br>Primers<br>Cp Mean (SD) | CIM Bennett<br>Primers<br>Cp Mean (SD) | P<br>value<sup>#</sup> | P<br>value<sup>+</sup> |
| --- | --- | --- | --- | --- | --- | --- |
| Overall | 107 | 33.1 (4.2) | 28.7 (4.3) | 27.1 (3.9) | <0.001 | 0.25 |
| Probable* | 84 | 33.1 (3.9) | 29.1 (4.0) | 27.6 (3.5) | <0.001 | 0.37 |
| Presumed* | 17 | 33.3 (5.4) | 27.8 (6.0) | 23.7 (4.9) | 0.037 | 0.73 |
| Possible* | 6 | 33.1 (6.6) | 25.6 (1.5) | 27.4 (6.2) | 0.44 | 1 |
| P value<sup>±</sup> |  | 0.99 | 0.48 | 0.12 |  |  |

\*As determined by two independent observers.
<sup>#</sup>Overall comparison of Cp values between swab and both CIM groups (One-way ANOVA).
<sup>+</sup>Compares CIM Dupuis Cp and CIM Bennett Cp.
<sup>±</sup>Overall comparison between observer groups

## Discussion

In suspected microbial keratitis, rapid and accurate diagnosis with immediate treatment are important to optimise clinical outcome. Although clinical features of HSK are essential to the diagnosis, reliance only on clinical features alone, may be misleading due to overlapping clinical findings caused by different conditions and infections, and excludes the ability to detect resistance mutations and or contact tracing.

Cell culture has been the traditional method for the detection of HSV-1, but has been largely replaced by PCR, due to its high sensitivity and shorter processing time. PCR has been optimised for the detection of HSV-1 from ophthalmic samples (12, 23-25) using swabs.(14) There is very little data, however, on the detection rates of HSV-1 from corneal and or conjunctival swabs in clinical practice and although anecdotal, many ophthalmologists do not collect samples in cases of suspected HSV-1 keratitis possibly due to the low yield and cumbersome nature of a swab. It is also unclear whether in clinical practice the majority of specimens are collected from the conjunctiva and tear film rather than from the corneal ulcer itself. It is not known whether a swab collects corneal epithelial cells, which is important for intracellular infections such as HSV-1. Impression cytology using a CIM, however, removes epithelial cells thus enabling the detection of intracellular microorganisms such as HSV-1. In addition, impression cytology has been shown to be useful for the diagnosis of a variety of infectious and non-infectious corneal conditions, including viral (15), fungal (26), acanthamoeba (27) and microbial keratitis (16), as well as for ocular surface neoplasia (17), keratoconjunctivitis sicca (18), vitamin A deficiency (28), and atopic keratoconjunctitivits.(29) In cases of suspected bacterial and acanthamoebic keratitis, a CIM has been shown to have higher sensitivity than more invasive methods and to be easy to use.(16) Importantly, the detection of human DNA extracted from corneal epithelial cells using a CIM and then amplified by the internal amplification control PCR, may be used as an indicator of sample quality.

HSV-1 also appears to be stable on CIM. Somerville et al. (30) recently demonstrated the stability of HSV-1 DNA recovery following inoculation of HSV-1 onto PTFE CIMs and storage at +4°C, −20°C and −70°C for up to 10 months. In this study, we obtained similar HSV-1 DNA Cp values to that demonstrated by Somerville et al. (30) both from CIMs extracted immediately following sample collection and those extracted 24 hours after collection and storage at +4°C. This suggests there is no significant reduction in HSV-1 DNA recovery should there be a 24 hour delay in sample processing. This is reflective of clinical settings in which samples often do not reach the laboratory until the following day after the sample has been collected.

We compared two sets of HSV-1 primers as one had been used for the testing of cases of encephalitis and the other had been optimised for the testing of eye swabs but no clinical sample testing was reported.(13) Both showed good and comparable sensitivity. In clinical practice, because topical anaesthetic is applied to the eye prior to a corneal sample (either a swab or a CIM) being collected, it was important therefore to demonstrate that there were no inhibitory effects on the PCR. Our *in vitro* data demonstrated good detection of HSV-1 DNA by PCR from CIM using both sets of primers with an end point of ≤10 virus genome copies/mL, which would be suitable for testing clinical samples.

A CIM is easy to use and may therefore be suitable for use by non-ophthalmologists or where less sophisticated biomicroscopes are available, such as in resource poor settings. In a recently published study (16) we compared the microbial detection rates of a corneal scrape to that using a CIM made from PTFE. The results using a CIM were significantly better than using a blade to detect bacteria and acanthamoeba from corneal ulcers in cases of suspected microbial keratitis.(16) The results of this study would suggest that a CIM may additionally be a simple and good alternative to using sharp instruments and swabs for the identification of HSV-1 in cases of suspected Herpes Simplex keratitis.

## References

1. FarooqAV, ShuklaD. 2012. Herpes simplex epithelial and stromal keratitis: an epidemiologic update. Survey of Ophthalmology 57:448–462.

2. Liesegang TJ. 1988. Epidemiology and natural history of ocular herpes simplex virus infection in Rochester, Minnesota, 1950-1982. Trans Am Ophthalmol Soc 86:688–724.

3. LabetoulleM, AuquierP, ConradH, CrochardA, DaniloskiM, BouéeS, Hasnaoui ElA, ColinJ. 2005. Incidence of herpes simplex virus keratitis in France. Ophthalmology 112:888–895.

4. GreenLK, Pavan-LangstonD. 2006. Herpes simplex ocular inflammatory disease. Int Ophthalmol Clin 46:27–37.

5. Farooq AV. 2017. Herpes Simplex Virus Keratitis and Resistance to Acyclovir. Cornea 36:e4–e5.

6. PiretJ, BoivinG. 2016. Antiviral resistance in herpes simplex virus and varicella-zoster virus infections: diagnosis and management. Curr Opin Infect Dis 29:654–662.

7. PanD, KayeSB, HopkinsM, KirwanR, HartIJ, CoenDM. 2014. Common and new acyclovir resistant herpes simplex virus-1 mutants causing bilateral recurrent herpetic keratitis in an immunocompetent patient. J Infect Dis 209:345–349.

8. KayeS, ChoudharyA. 2006. Herpes simplex keratitis. Progress in Retinal and Eye Research 25:355–380.

9. SubhanS, JoseRJ, DuggiralaA, HariR, KrishnaP, ReddyS, SharmaS. 2004. Diagnosis of herpes simplex virus-1 keratitis: comparison of Giemsa stain, immunofluorescence assay and polymerase chain reaction. Curr Eye Res 29:209–213.

10. MengelleC, Sandres-SaunéK, MiédougéM, MansuyJM, BouquiesC, IzopetJ. 2004. Use of two real-time polymerase chain reactions (PCRs) to detect herpes simplex type 1 and 2-DNA after automated extraction of nucleic acid. J Med Virol 74:459–462.

11. CoreyL, HuangM-L, SelkeS, WaldA. 2005. Differentiation of herpes simplex virus types 1 and 2 in clinical samples by a real-time taqman PCR assay. J Med Virol 76:350–355.

12. RyncarzAJ, GoddardJ, WaldA, HuangML, RoizmanB, CoreyL. 1999. Development of a high-throughput quantitative assay for detecting herpes simplex virus DNA in clinical samples. Journal of Clinical Microbiology 37:1941–1947.

13. DupuisM, HullR, WangH, NattanmaiS, GlasheenB, FuscoH, DziguaL, MarkeyK, TavakoliNP. 2011. Molecular detection of viral causes of encephalitis and meningitis in New York State. J Med Virol 83:2172–2181.

14. BennettS, CarmanWF, GunsonRN. 2013. The development of a multiplex real-time PCR for the detection of herpes simplex virus 1 and 2, varizella zoster virus, adenovirus and Chlamydia trachomatis from eye swabs. J Virol Methods 189:143–147.

15. ThielMA, BossartW, BernauerW. 1997. Improved impression cytology techniques for the immunopathological diagnosis of superficial viral infections. British Journal of Ophthalmology 81:984–988.

16. KayeS, SuekeH, RomanoV, ChenJY, CarntN, TuftS, NealT. 2015. Impression membrane for the diagnosis of microbial keratitis. Br J Ophthalmol 100:bjophthalmol– 2015–307091–610.

17. BarrosJ de N, AlmeidaSRA de, LowenMS, CunhaMCD, GomesJÁP. 2015. Impression cytology in the evaluation of ocular surface tumors: review article. Arq Bras Oftalmol 78:126–132.

18. LopinE, DeveneyT, AsbellPA. 2009. Impression cytology: recent advances and applications in dry eye disease. Ocular Surface 7:93–110.

19. ChenC-Y, ChiKH, AlexanderS, IsonCA, BallardRC. 2008. A real-time quadriplex PCR assay for the diagnosis of rectal lymphogranuloma venereum and non-lymphogranuloma venereum Chlamydia trachomatis infections. Sexually Transmitted Infections 84:273–276.

20. JonesCD, YeungC, ZehnderJL. 2003. Comprehensive validation of a real-time quantitative bcr-abl assay for clinical laboratory use. Am J Clin Pathol 120:42–48.

21. GoldschmidtP, RostaneH, Saint-JeanC, BatellierL, AlouchC, ZitoE, BourcierT, LarocheL, ChaumeilC. 2006. Effects of topical anaesthetics and fluorescein on the real-time PCR used for the diagnosis of Herpesviruses and Acanthamoeba keratitis. British Journal of Ophthalmology 90:1354–1356.

22. LiJ, FineJ. 2004. On sample size for sensitivity and specificity in prospective diagnostic accuracy studies. Stat Med 23:2537–2550.

23. BispoPJM, DavoudiS, SahmML, RenA, MillerJ, RomanoJ, SobrinL, GilmoreMS. 2018. Rapid Detection and Identification of Uveitis Pathogens by Qualitative Multiplex Real-Time PCR. Invest Ophthalmol Vis Sci 59:582–589.

24. DominguezSR, PrettyK, HengartnerR, RobinsonCC. 2018. Comparison of Herpes Simplex Virus PCR with Culture for Virus Detection in Multisource Surface Swab Specimens from Neonates. Journal of Clinical Microbiology JCM.00632–18.

25. YoshidaM, HariyaT, YokokuraS, MaruyamaK, SatoK, SugitaS, TomaruY, ShimizuN, NakazawaT. 2018. Diagnosing superinfection keratitis with multiplex polymerase chain reaction. J Infect Chemother.

26. JainAK, BansalR, FelcidaV, RajwanshiA. 2007. Evaluation of impression smear in the diagnosis of fungal keratitis. Indian J Ophthalmol 55:33–36.

27. SawadaY, YuanC, HuangAJW. 2004. Impression cytology in the diagnosis of acanthamoeba keratitis with surface involvement. AJOPHT 137:323–328.

28. KhanAN, HudaS, AhmedAN, HossainT, SultanaN, AliSM. 1992. Detection of early xerophthalmia by impression cytology and rose Bengal staining--a comparative study. Bangladesh Med Res Counc Bull 18:1–11.

29. AragonaP, RomeoGF, PuzzoloD, MicaliA, FerreriG. 1996. Impression cytology of the conjunctival epithelium in patients with vernal conjunctivitis. Eye 10 (Pt 1):82–85.

30. SomervilleTF, CorlessCE, NealT, KayeSB. 2018. Effect of storage time and temperature on the detection of Pseudomonas aeruginosa, Acanthamoeba and Herpes Simplex Virus from corneal impression membranes. J Med Microbiol.

